# Atlas of lysine acetylation in the mouse

**DOI:** 10.64898/2026.01.09.698739

**Authors:** Ross W. Soens, Benton J. Anderson, Noah M. Lancaster, Mukesh Kumar, Julia K. Hanssen, Andrea Galmozzi, Timothy Grant, Katherine A. Overmyer, Joshua J. Coon

## Abstract

Lysine acetylation has widespread ramifications from genetic regulatory effects to modulation of enzymatic function. With improved acetyl-lysine enrichment technologies and advances in mass spectrometer speed and sensitivity, we present a comprehensive atlas of the mouse acetylome comprising 17,952 unique lysine acetylation sites across 4,340 proteins and 15 tissues. This resource, which nearly doubles the known mouse acetyl-lysine catalog, shows at least 14% of the acetylome is shared across tissues. We focus our investigation on several acetylated proteins, including ribosomal acetylation and its potential to extend ribosomal half-life in the liver and pancreas. Additionally, we identify a novel acetylation event in the active site of carnitine O-acetyltransferase (Crat) that also mirrors tissue-specific Crat activity. By integrating these data with human pathogenic variants, we identify acetyl-lysine residues on cardiac troponin and homogentisate dioxygenase that likely mimic disease-causing mutations. This resource provides a foundational framework for investigating protein acetylation in metabolic health and disease.

## Introduction

Post-translational modifications provide an additional layer of complexity upon the central dogma, allowing cells to adjust the utility of proteins well past what their initial coding and alternative splicing allows. Due to the multitude of protein functions that involve electrostatic interactions, PTMs that add or remove a charge from a residue’s side chain provide a key molecular toolset. One such prominent modification can be found in the acetylation of lysine, whereby an acetyl group from acetyl-coenzyme A (acetyl-CoA) is transferred to the primary amine group thus neutralizing its positive charge at physiologic pH^1^. Originally identified in its role regulating the function of histones^2,3^, mass spectrometry (MS)-based proteomics has provided evidence that lysine acetylation is widespread across the proteome^4^. This reversible transfer occurs both enzymatically and nonenzymatically at varying levels of stoichiometry based on subcellular localization, the surrounding chemical environment, and the activity of acetyltransferases/deacetylases. For example, due to the high abundance of acetyl-CoA and high pH, mitochondrial proteins are more commonly acetylated than phosphorylated^5,6^. These acetyl-lysine sites provide key regulatory control over the bioenergetic capacity of mitochondria^7,8^.

Lysine acetylation sites are implicated in numerous cellular processes including DNA damage repair, metabolic control, transcription, and translation^9^. In addition, acetyl-lysine residues are associated with disease, e.g., acetyl-lysine modifications of Tau impact its aggregation in Alzheimer’s^10–12^ and mutations of lysine acetyltransferase/deacetylases can promote malignancy or act as tumor suppressors in various cancers^13,14^. Along with the long-appreciated role of lysine acetylation regulating the DNA binding capacity of histones, acetyl-lysine residues have been found to play a major role in metabolic regulation including but not limited to glycolysis, gluconeogenesis, the TCA cycle, the urea cycle, and fatty acid metabolism^15,16^.

Despite its widespread prevalence and biological importance, lysine acetylation remains understudied owing primarily to the difficulty of its detection. Like many other PTMs, lysine acetylation occurs at low stoichiometry such that standard global proteome measurements rarely detect it. And, unlike phosphorylation where simple metal-affinity chromatographic enrichment technologies permit routine enrichment and detection of tens of thousands of sites per hour, enrichment of acetylated lysine residues is more nuanced and relies on antibody-based technologies^17–19^. Another difficulty in studying lysine acetylation is its relatively low stoichiometry – estimated at ∼ 0.02%^20,21^. Given these low levels, even methods with optimized enrichment strategies likely miss a large subset of acetyl-lysine containing peptides whose overall abundance falls below the mass spectrometer system’s limit of detection^22^.

The most extensive acetyl-lysine mapping effort by Lundy et. al. described ∼15,000 lysine acetyl sites across 16 rat tissues. This 2012 study leveraged the best technology at the time, yet in the decade since, technologies have rapidly improved enrichment yields and mass spectrometry sensitivity and speed. Specifically, we leveraged newly developed antibody enrichment technologies^18^ and the Orbitrap Astral Zoom hybrid mass spectrometer^23^ with data independent acquisition to generate the most extensive atlas of mammalian lysine acetylation to date. We provide evidence for 17,952 acetyl-lysine sites across 4,340 proteins in less than two days of MS analysis. This atlas, compiled into a public resource, highlights lysine acetylation as more universal across tissues than previously described^8^. We further expand our understanding of the potential for acetylation to regulate key cellular mechanisms, such as ribosomal recycling and mitochondrial acetyl-CoA level maintenance.

## Results

### Improved characterization of the lysine acetylproteome

To update the observable mouse lysine acetylome, we first had to establish methods for consistent acetyl-lysine peptide sample preparation and analysis (**Figure 1A**). For initial method development, acetyl-lysine peptides were enriched from a tryptic digest of mouse liver proteins. We hypothesized that recent improvements in mass spectrometry instrumentation could enhance detection of these low abundance peptides. Specifically, we recently described the use of the Orbitrap Astral system and its ability to collect high sensitivity MS/MS scans at ∼ 200 Hz^17^. When coupled with data independent acquisition (DIA) methods this platform allowed for the detection of ∼ 30,000 unique phosphorylation sites within a single 15-minute separation^17^. Since that time, an even faster implementation of the Astral system (Orbitrap Astral Zoom) has boosted scan rates to ∼ 270 Hz ^23^. To evaluate the ability of this instrument to detect acetyl-lysine peptides we performed nanoflow liquid chromatography tandem mass spectrometry (nLC-MS/MS) on enriched acetyl-lysine peptides (250 ng load). To ensure optimal settings, we varied numerous parameters including: sample injection time, precursor mass-to-charge (*m/z*) range, DIA window width, MS/MS AGC target, HCD collision energy, gradient length, and RF lens settings (**Supplementary Figure 1 A-G**). To support further validation of acetyl-lysine peptides, the MS/MS scan range was lowered to 140 *m/z* to allow for detection of immonium ions, a known reporter ion for acetyl-lysine containing precursors^24^ (**Supplementary Figure 1H**). We also evaluated offline high-pH reverse phase fractionation to boost identifications; however, due to the low mass yield of acetyl-enriched samples (∼40µg/4mg liver peptides), fractionation did not assist in better acetyl-site identification (**Supplementary Figure 1I**).

**Figure 1.**
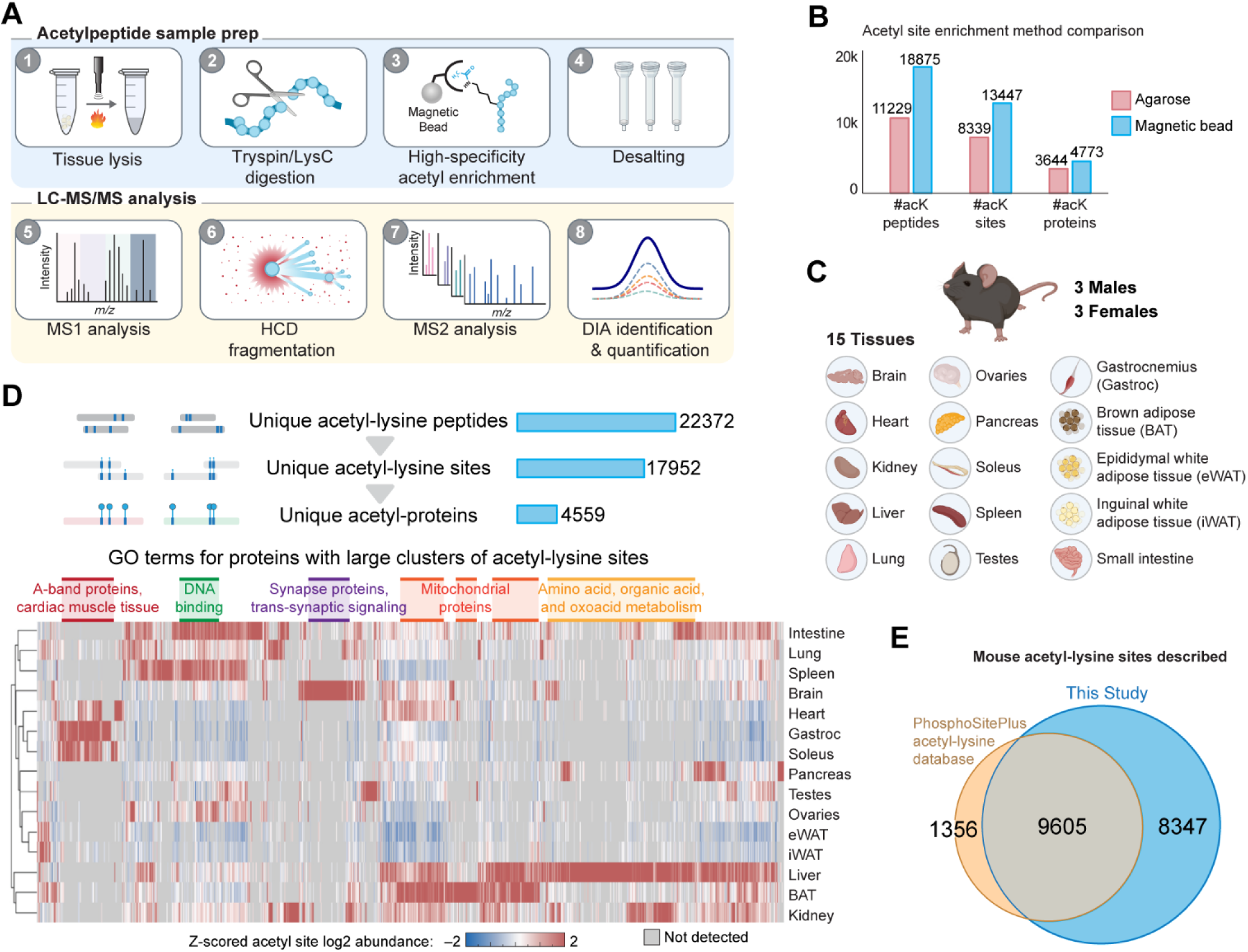
Acetyl-lysine atlas workflow and key features. **A** Mouse acetyl-lysine atlas workflow. **B** Comparison of acetyl-lysine sites, peptides, and proteins identified using either agarose (PTMScan Classic) or magnetic bead based high specificity (PTMScan HS) enrichment protocols (n=1, mouse liver). **C** Mouse acetyl-lysine atlas tissue overview. **D** Clustered heat map of Z-scores of all observed acetyl-lysine site log2 abundances with unique groups and broad atlas overview above. Note selected GO terms highlighted over columns show enriched acetyl-lysine pathways. **E** Venn diagram of all detected mouse acetyl-lysine sites in our study compared to the PhosphositePlus database.

In previous acetyl-lysine mapping efforts, enrichment techniques utilized anti-acetyl-lysine antibodies conjugated to agarose beads^8,21,25^. However, modern immunoenrichment strategies have gravitated towards magnetic-bead based approaches for more streamlined and effective workflows^26–29^. We recently developed a magnetic-bead based enrichment strategy, the High Specificity (HS) PTMScan acetyl-lysine immunoenrichment kit (PTMScan HS), which crosslinks the anti-acetyl-lysine antibodies to magnetic beads and incorporates updated binding and washing buffers from the original agarose-bead based protocol (PTMScan acetyl-lysine). These modifications inhibit co-elution of the antibody with the eluted acetyl-peptides and increase specificity of enrichment, leading to a two to three-fold improved detection capability according to a recent publication^18^. We similarly evaluated the HS-PTMScan technology by enriching acetyl-lysine peptides from mouse liver and found a 1.6-fold increase in detected acetyl-lysine sites compared to the classic PTMScan method (**Figure 1B**). With the combination of the improved acetyl-lysine enrichment and the optimized Astral Zoom acquisition, we detected 13,447 unique acetyl-lysine sites, from over 18,000 acetyl-lysine peptides in a single-shot, 30 minute LC-MS/MS method (**Figure 1B**).

### Construction of a comprehensive atlas of the acetyl-lysine proteome

To establish the comprehensive mouse acetyl-lysine atlas, we enriched acetyl-lysine peptides from fifteen unique mouse tissues collected from male and female C57BL6/J mice. **Figure 1C** details the collected tissues: pancreas, small intestine, spleen, liver, kidney, heart, lung, gastrocnemius muscle, soleus muscle, brain, brown adipose tissue (BAT), epididymal white adipose tissue (WAT), inguinal WAT, testes, and ovaries from mice aged 47 days. Briefly, each tissue was pooled by sex, then protein was extracted, digested with trypsin/LysC, and enriched for acetyl-lysine containing peptides using the PTMscan HS enrichment kit. Both enriched and non-enriched peptides were separately analyzed in triplicate with the optimized nLC-MS/MS single shot method detailed above, yielding 84 raw data files.

Altogether these experiments yielded 17,938 unique acetyl-lysine sites across 4,559 proteins, many of which exhibit tissue specificity and associate with tissue function (**Figure 1D**). For example, A-band proteins have acetyl-lysine signatures that are common across muscle tissues gastrocnemius, soleus, and heart. In contrast, acetylation sites on synapse-associated proteins are uniquely enriched in brain tissue. Further, proteins involved in amino acid and organic acid metabolism have an enriched acetylation signature in the liver. To assess the discovery potential of our new atlas, we cross-referenced our results with the PhosphoSitePlus (PSP) database^30^, which compiles acetyl-lysine sites, among other PTMs. The overlap of our experimental results with the reference PhosphoSitePlus database was 9,605 acetyl-sites, ∼90% of the database. However, with evidence for an additional 8,347 unique acetyl sites, this work nearly doubles the current PhophoSitePlus catalog of known mouse lysine acetylation sites (**Figure 1E**). Finally, we supply these data as a resource for the community (**Supplemental Data 1**).

### The mouse acetyl-atlas offers unique insight into tissue-specific lysine acetylation

This extensive multi-tissue acetyl-proteome analysis provides a new benchmark for comprehensive acetylomics and provides a new resource for the community. This atlas builds on the pioneering work of Lundy et al. who identified 15,474 acetyl-lysine sites across sixteen rat tissues (**Figure 2A**)^8^. This study identifies 17,938 acetyl-lysine sites in total, approximately 16% more, and shows a marked increase in the number of acetyl-lysine sites per tissue, suggesting our updated methodology offers improved reproducibility and lower limits of detection. This trend is also evident in the protein level data where both studies identify a similar number of acetyl-proteins (4,559 versus 4,247), our study on average doubles the number of acetyl-proteins detected per tissue, where among tissues analyzed in both studies, we detect an average of 3,013 acetylproteins per tissue, versus 1,430 per tissue in Lundby (**Figure 2B**). The largest group of acetyl-lysine containing proteins have only one acetylated residue (**Figure 2C**). Globally, the average site/protein is 4.13 with a median of 2 sites/protein. However, many outliers exist, such as the 626 acetyl-lysine sites found on Titin or the high rate of lysine acetylation (10 out of 12 lysines) on ATP synthase peripheral stalk subunit d (Atp5pd).

**Figure 2.**
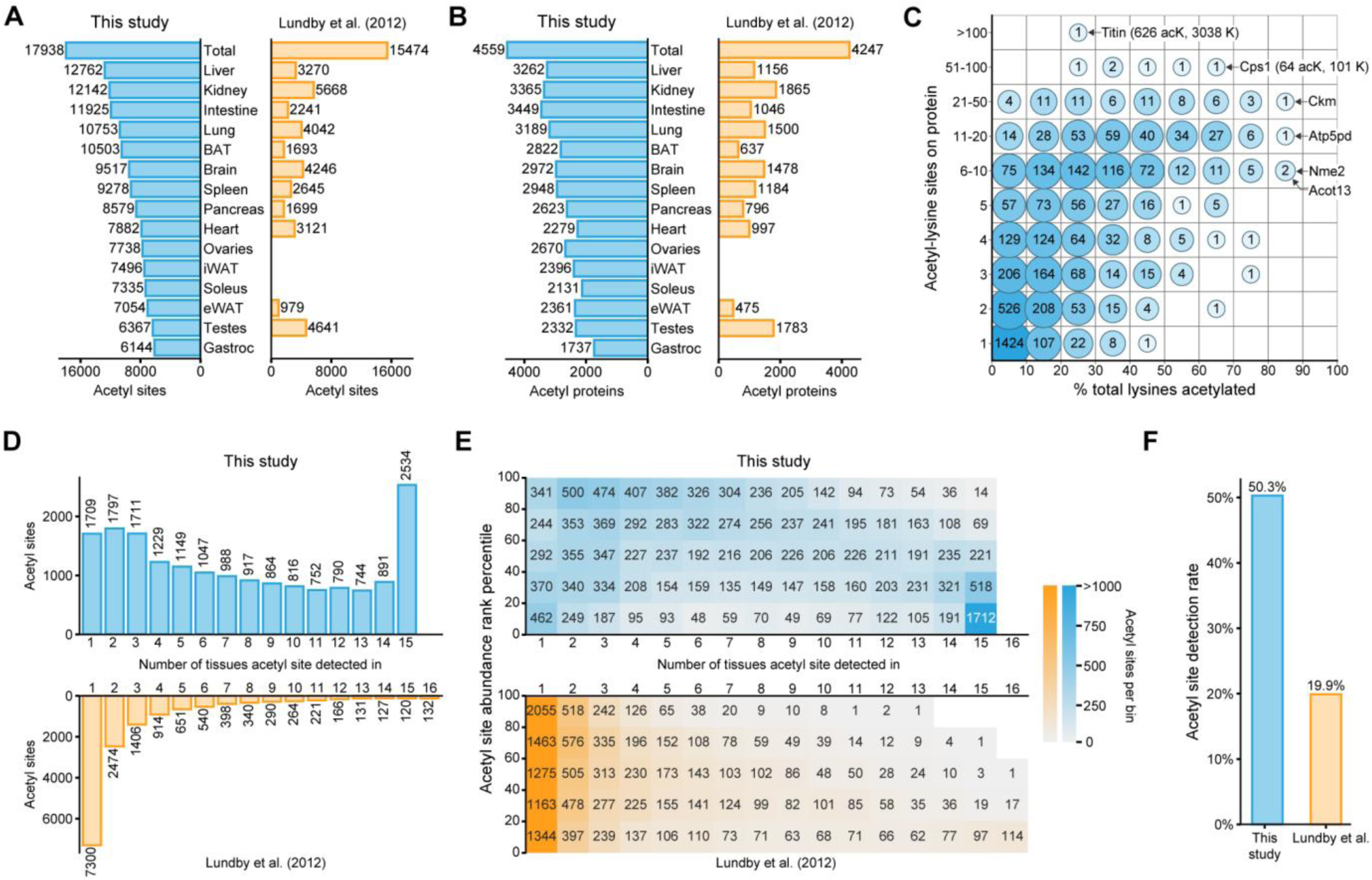
Acetyl-lysine atlas tissue distribution. **A** Number of acetyl-lysine sites detected by tissue and compared to data of Lundby et al. **B** Number of acetyl-lysine modified proteins by tissue and compared with Lundby et al^8^. **C** Map of acetyl-lysine sites (detected in this study) per protein across all tissues against the total percentage of lysines acetylated per protein. **D** Tissue specificity of detected acetyl-lysine sites compared with Lundby et al. **E** Comparison of mean acetyl site abundanc e versus tissue detection rate, in this work (upper) and Lundby et al. (lower). **F** Acetyl site detection rate in this work versus Lundby et al.

To examine tissue specificity of lysine acetylation, we plotted the distribution of sites found across tissues (**Figure 2D**). When compared to the previous study, we observe a strikingly higher number of sites detected across all tissues. While the largest group (14%) of acetyl sites is found in every tissue, we find 20% of sites in only one or two tissues. This stands in contrast to Lundby et al. who found that acetylation is sparsely distributed across tissues with 47% of all acetyl sites only detected in one tissue. To understand whether quantitative trends associate with the difference in detection rates between these studies, we assessed the acetyl site mean quantitation versus the number of tissues a site was detected in (**Figure 2E**). With our approach, we observed high tissue overlap even with low-abundance proteins, suggesting that higher sensitivity methods and DIA approach offered more data completeness. Our work redefines the coverage of acetyl-lysines across tissues from being one of relatively high sparsity with less than 20% completeness, to a more highly conserved PTM with on average 50% detection across tissues (**Figure 2F**).

### Lysine acetylation is associated with tissue-specific function and protein structural features

To begin to dive deeper into global analysis, we started with an unsupervised approach via principal component analysis (PCA) of all unique acetyl-lysine sites (**Figure 3A**). Here, we note that tissues group on principal components one and two (PC1 and PC2) by function, such as muscles (gastrocnemius, soleus, heart) and white adipose tissues (eWAT, iWAT). Differences between acetyl sites between sexes exist but were much smaller than those between tissues. Based on the loadings, the top drivers of PC1 were primarily acetyl-lysine sites on metabolic and muscle proteins (**Supplementary Figure 2**). Among the top drivers were heat shock proteins, an observation that supports previous studies finding evidence of differential expression of heat shock proteins based on sex and tissue of origin^31–33^. The top drivers of PC2 were populated by acetyl-lysine sites on core mitochondrial metabolic enzymes and histone acetyltransferases (**Supplementary Figure 2**). Accordingly, PC2 appears to separate high metabolic activity tissues (BAT and muscle) compared to lower metabolic activity tissues like ovaries and spleen. This distribution could be attributed to the higher acetyl-CoA levels found in mitochondria^34^, specifically those mitochondria of highly metabolically active tissues^35^. Together, these observations indicate this atlas contains a wealth of information regarding tissue specific differences in lysine acetylation.

**Figure 3.**
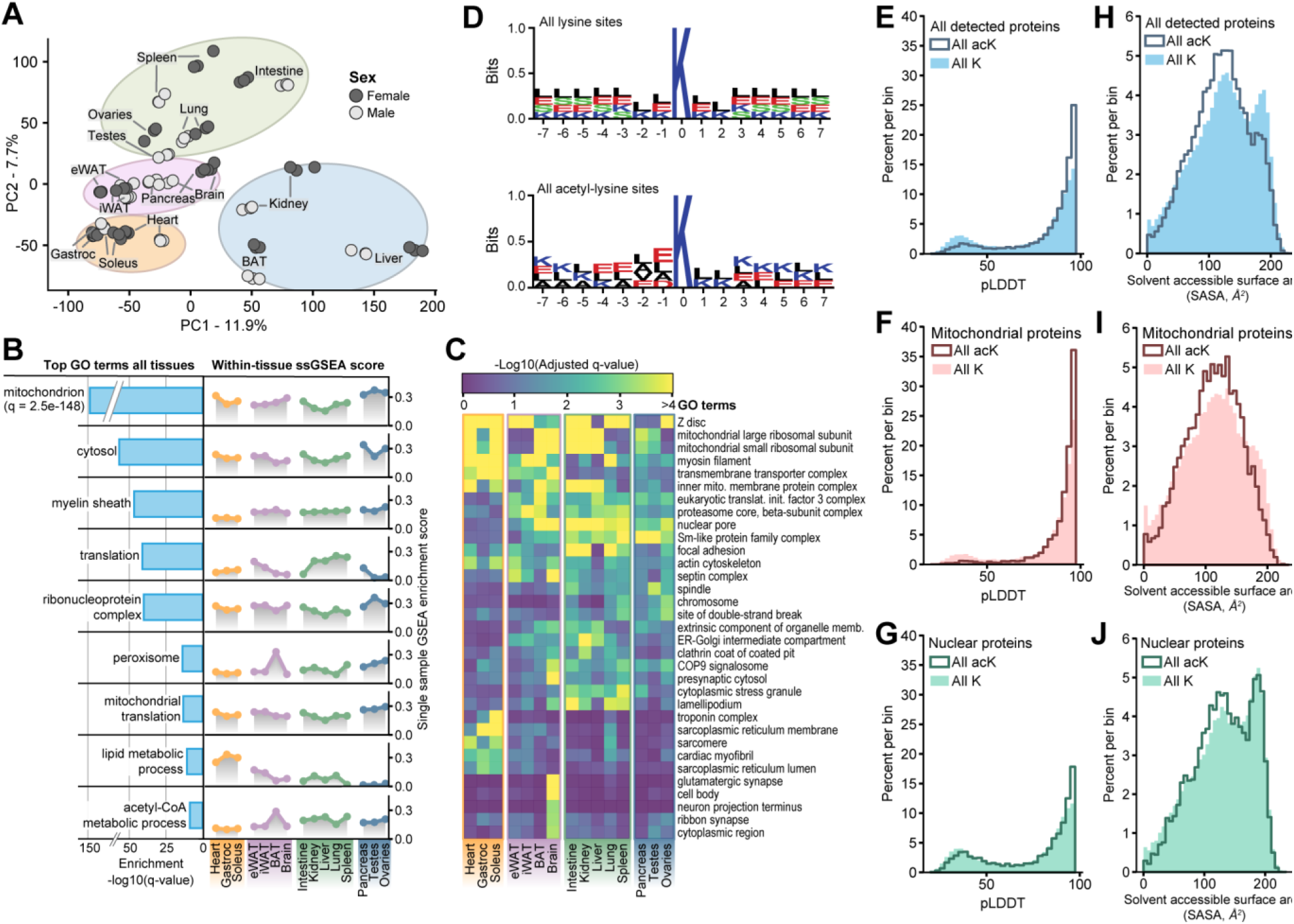
Tissue grouping, GO term analysis, sequence, and structural specificity of acetyl-lysine sites. **A** Principal Component Analysis (PCA) of all acetyl-lysine sites detected across all tissues, with sex indicated by color. **B** Global gene ontology (GO) term enrichment analysis relative to all detected proteins across all tissues (left). GSEA scores plotted for each tissue and each GO term (right). **C** Clustered heatmap showing GO terms with unique acetyl-lysine enrichment profiles across tissues. **D** Sequence logo plot for all lysines throughout the mouse FASTA (top) or all detected acetyl-lysine sites (bottom). **E** Per residue model confidence (pLDDT) distribution of all lysine sites across the mouse FASTA, all detected acetyl-lysine sites, or all acetyl sites detected in **F** mitochondria or **G** the nucleus. **H** Solvent accessible surface area distribution of all lysine sites across the mouse FASTA, all detected acetyl-lysine sites, or all acetyl sites detected in **I** mitochondria or **J** the nucleus.

Using acetyl-lysine sites for enrichment analysis^36,37^, we find that the mitochondrion gene ontology (GO) term is the most significantly enriched category; other top GO terms include cytosol, amino acid metabolism, and translation, etc. (**Figure 3B**). From this global enrichment, we further evaluated individual tissue enrichments for each of these categories^38^. For mitochondrion, we found that liver, kidney, and BAT exhibited the greatest magnitude of enrichment. In addition, we found other expected trends, like brain having the highest magnitude of enrichment for the myelin sheath GO term. Interestingly we found that for ribonucleoprotein complex and translation GO terms, the pancreas and liver were the most elevated. A breakdown of the top ten enriched GO terms of acetyl-lysine sites within each individual tissue can be found in **Supplementary Figure 3A**. As observed in past work^8^, lysine acetylation sites occur on proteins that also have tissue specific functions.

In contrast to the global GO term enrichment analysis, we extracted GO terms that exhibited high variance across tissues (**Figure 3C**). We highlight 33 of these GO terms; for example, neuron and synapse-specific protein acetylation drives the unique profile of brain tissue^39^ while muscle-specific protein acetylation, such as sarcomere and troponin protein lysine acetylation, defines a unique acetyl-enrichment group in the heart, gastrocnemius, and the soleus^40,41^. We also observe unique lack of enrichment in otherwise commonly acetylated proteins, such as Z disc proteins^42^ with lower lysine acetylation in the pancreas, liver, and testes. A complete table of GO term enrichment results and ssGSEA scores is given in **Supplemental Data 2.**

Previous studies have determined that acetyl-lysine sites are often flanked by one or multiple other lysine residues, with a noted increase in negatively charged residues directly upstream ^8,43^. With our expanded compendium of lysine acetylation sites, we first calculated the sequence motif of all lysines, modified or unmodified, and compared that to the motif of acetyl-lysine sites (**Figure 3D**). With this analysis we confirmed and offered supporting evidence that acetyl-lysines occur directly downstream of negatively charged residues and are often surrounded by other lysine residues^8,44^. We observe no notable differences across tissuesor subcellular localizations (**Supplementary Figure 3B**). This is in contrast to Lundby et. al., who found subcellular-specific motifs^8^. We suspect that this observation could have been due to lower site coverage.

A limitation of traditional primary sequence-based motif mapping is that it does not consider secondary and tertiary structural information, which doubtless is a key component of lysine acetylation dynamics. To address this shortcoming, we leveraged AlphaFold2 structural predictions to access the three-dimensional environments of all detected acetyl-lysine sites^45^. These predicted structures have varying confidence levels, which are calculated at a single amino acid resolution (i.e., per residue model confidence score, pLDDT). These confidence scores are associated with the flexibility of a protein structure, with low confidence levels being associated with more disordered regions. **Figure 3E** plots the distribution of pLDDT scores for acetylated lysines and all lysines. Note the striking shift of acetylated lysine residues toward higher model confidence values, suggesting these sites exist in more highly structured regions. This observation is counter to what has been previously reported for S/T/Y phosphorylation, where the modification is enriched in disordered regions ^17,46,47^. To determine whether cellular compartments have differences in lysine-acetylation structural preferences we also plotted the score distributions for nuclear and mitochondrial acetyl-lysine sites (**Figure 3F, G**). Interestingly, mitochondrial proteins show even stronger preference for ordered structural regions whereas the acetylated-lysines on nuclear proteins have a score distribution that more closely tracks with non-acetylated lysine residues. This discrepancy could be attributed to high levels of acetyltransferase activity in the nucleus compared to more prevalent non-enzymatic acetylation in mitochondria^48^.

Our observation that lysine-acetylation is enriched in structured regions motivated us to investigate other structural features. Specifically, we calculated the solvent accessible surface area (SASA)^49–51^ score of each residue, which can act as a proxy for the position of the residue relative to the interior (less solvent accessible) or exterior (more solvent accessible) of the protein. Plotting these SASA scores (**Figure 3H**) revealed a bimodal distribution for all lysines within the mouse; however, acetyl-lysines were skewed towards lower solvent accessibility. This trend was even more prevalent within mitochondrial acetyl-lysine sites but for nuclear-localized sites the distribution more closely matched that of all lysine residues (**Figure 3I, J**). We postulate that this stark difference is due to the differential primary method of lysine acetylation across the cell. Enzymatic acetylation, for example by nuclear histone acetyltransferases, would naturally require site access and would be inhibited for buried residues. On the other hand, non-enzymatic lysine acetylation would have access to sites within folded pockets that neither acetyltransferases nor deacetylases could reliably act upon. Over time, proteins with longer half-lives, such as those found within mitochondria^52^, would gradually build up internal lysine acetylation, leading to a survivorship bias defining the distributional skew in acetyl-lysine solvent accessible area we have observed. Due to the higher concentration of acetyl-CoA^34^, and the preference for lower solvent accessibility, these data support previous observations that the primary mode of mitochondrial lysine acetylation is likely non-enzymatic^53,54^.

### Ribosomal protein lysine-acetylation and implications for homeostasis

Ribosomal subunits are among the most highly and consistently acetylated proteins across all measured tissues (i.e., 360 acetyl-lysine sites on 70 ribosomal subunits). Ribosomal protein acetylation is essential for maintaining proper ribosomal assembly and function^55^. **Figure 4A** presents an overview of global ribosomal lysine acetylation in the mouse – note the especially elevated levels in liver and pancreas. **Figure 4B** presents a three-dimensional model of the ribosome where these sites are evenly and broadly distributed across the entire complex. A sequence motif analysis of all lysine residues in the ribosome highlights the abundance of basic, positively charged, residues in the ribosome whereas acetyl-lysines tend to be downstream of acidic, negatively charged, residues (**Supplementary Figure 4A**). This observation is consistent with the known function of ribosomal structural proteins in stabilizing the interactions with negatively charged rRNA backbones.

**Figure 4.**
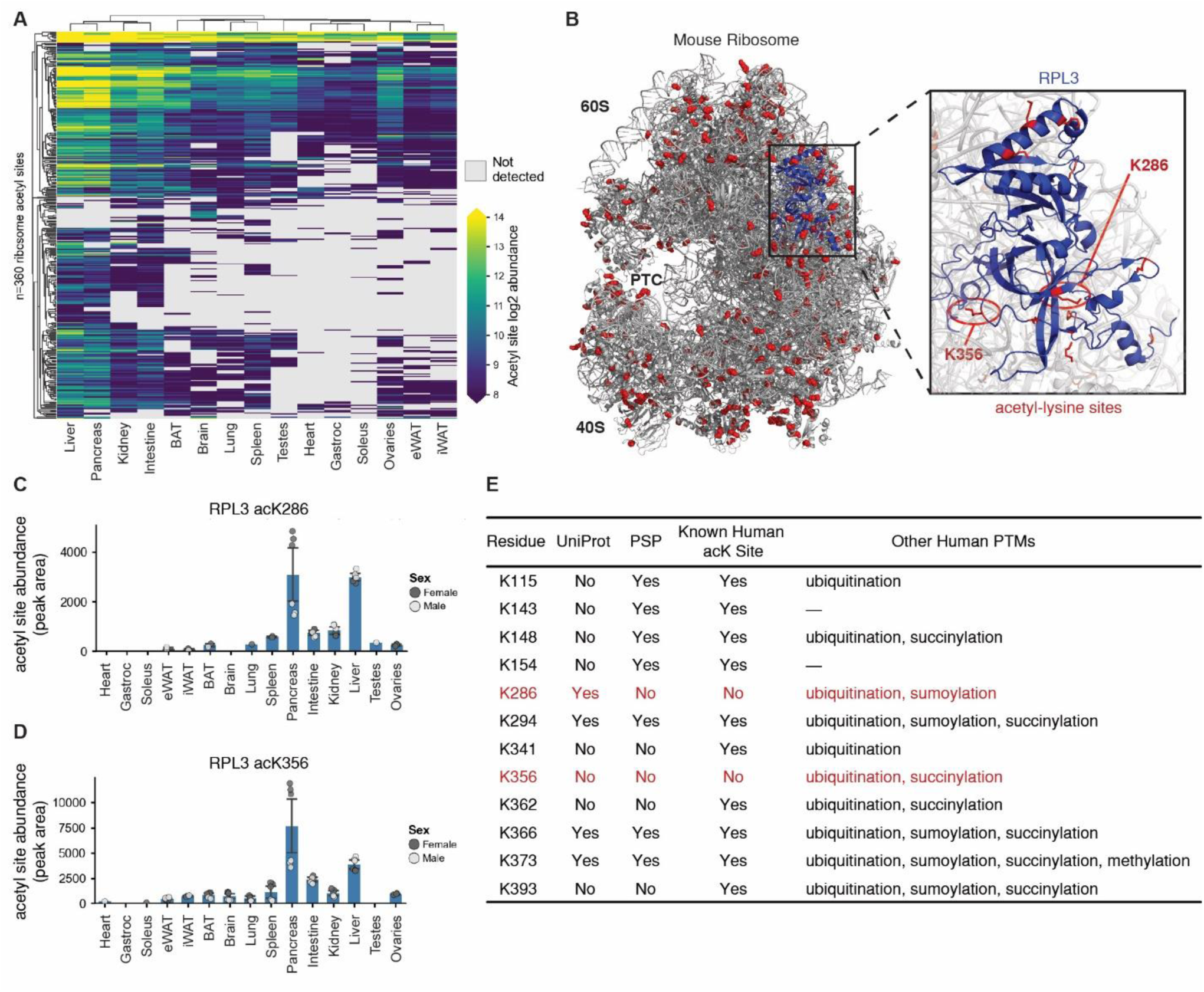
Ribosomal lysine acetylation coverage. **A** Clustered heatmap plotting all detected ribosomal protein-specific lysine acetylation abundances. **B** Mouse ribosome cryo-EM structure. Detected lysine acetylation sites colored in red. RPL3 colored in blue. PTC: peptidyl transferase center, RCSB PDB: 7CPU. **C** Abundance of RPL3 acK286 across tissues. **D** Abundance of RPL3 acK356 across tissues. **E** Table showing all detected RPL3 lysine acetylation sites with corresponding presence/absence in the UniProt or PSP database, with known human acetyl site listed as well as other known human PTMs.

Highlighted in **Figure 4B** is the subunit RPL3, one of the most acetylated ribosomal proteins observed, containing twelve unique lysine acetylation sites. RPL3 functions as a structural component of the large 60S ribosomal subunit opposite the peptidyl transferase center (PTC)^56^. Four of the twelve mouse sites are novel, i.e., not yet annotated in UniProt or PSP (**Figure 4E**). Two acetyl-lysine sites have not been analogously observed in PSP human studies, acK286 and acK356. Both sites are present at varying levels across all tissues but entirely absent in muscles (**Figure 4C, D and Supplementary Figure 4C**). Ten of these RPL3 acetyl-lysine sites are known human ubiquitination sites (**Figure 4E**). Because acetylation of lysine would prohibit ubiquitination^57^, these sites could therefore be used as a mechanism for increasing ribosomal half-life through inhibition of the ubiquitin-proteasome protein degradation pathway.

Elevated ribosomal lysine acetylation in the pancreas could support sustained translational activity during the post-prandial state, enabling the pancreas to produce insulin and reduce blood glucose (**Supplementary Table 1**). Reduced acetylation levels of such sites in muscles could be due to the high protein turnover rates found within these tissues^58–60^. This postulated role of acetylation elongating ribosomal half-life would require further study.

### Identification of novel lysine acetylation in Crat active site

Owing to the central role of mitochondria in acetyl-CoA production and cellular metabolism, this organelle exhibits the highest levels of lysine acetylation across the cell^6,48,61–64^. Our study both confirms and expands on this observation; for example, the top ten drivers of PC2 (**Figure 3A, Supplementary Figure 2C**) are dominated by mitochondrial enzymes. Further, the top GO hits across all tissues are mitochondrial (**Figure 3B**). And, given the observation that the mitochondrial acetyl-lysine site distribution is weighted towards less solvent accessible residues (**Figure 3G**), we supposed our dataset contained novel and potentially interesting acetyl-lysine sites within catalytic pockets. To investigate, we plotted all 5,728 mitochondrial lysine acetylation sites and their relative abundances across tissues (**Figure 5A**). These data showed that mitochondrial lysine acetylation was most abundant in brown adipose tissue, kidney, and liver. As previously suggested, these tissues are highly metabolically active, produce more acetyl-CoA, and consequently have elevated lysine acetylation^65^. Sequence motifs of mitochondrial acetyl-lysine residues showed no discernable difference from global acetyl-lysine motifs (**Supplementary Figure 5A**).

**Figure 5.**
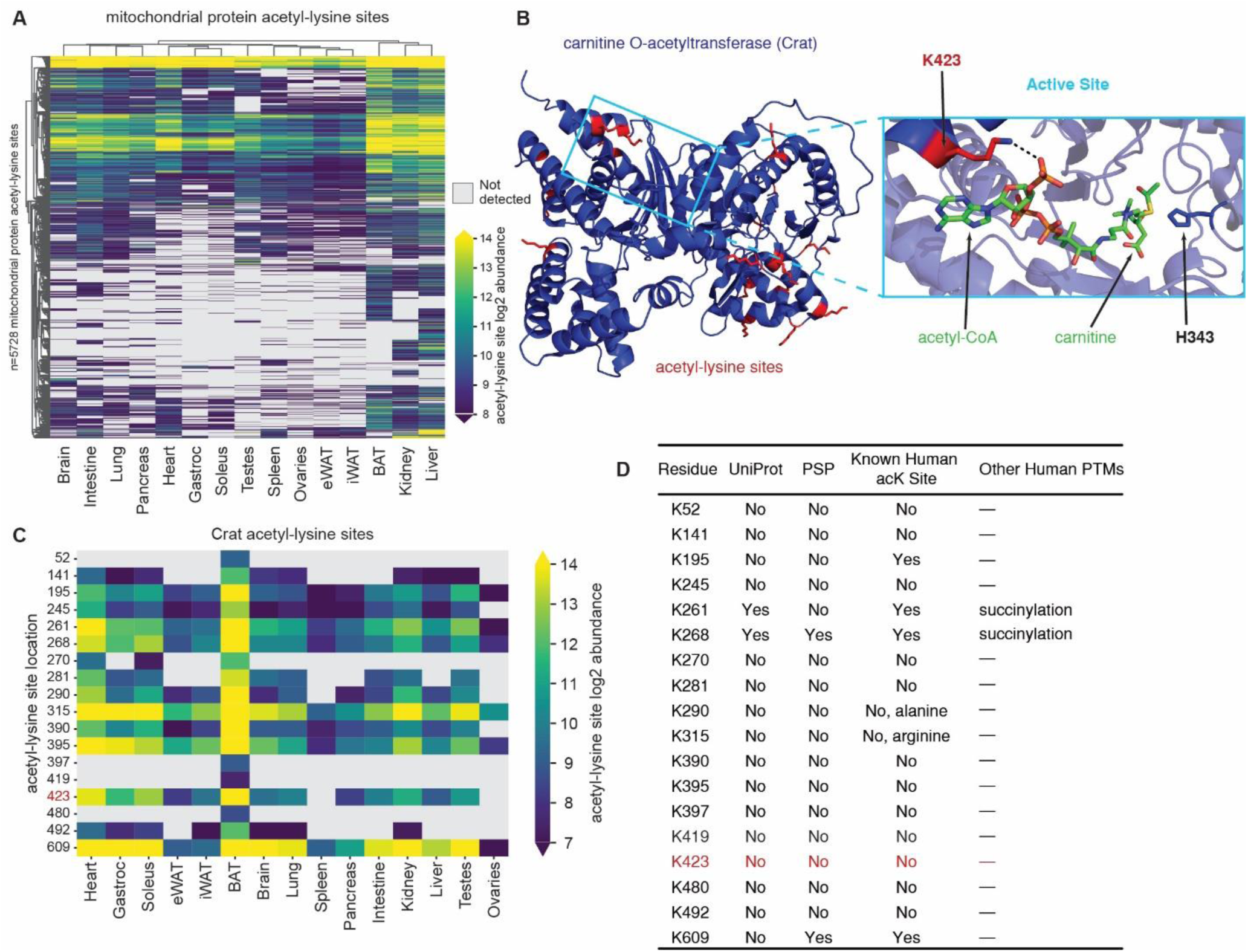
Mitochondrial lysine acetylation and Crat acetylation. **A** Clustered heatmap plotting all detected acetyl-lysine abundances on mitochondrial proteins. **B** Mouse Crat crystal structure. Detected lysine acetylation sites colored in red. Crat colored in blue. Active site substrate atoms colored as follows; carbon:green, nitrogen: blue hydrogen/lysine carbons: red, phosphate: orange, sulfur: yellow, active site histidine: dark blue. RCSB PDB: 2H3P. **C** Clustered heatmap of all identified Crat lysine acetylation sites. **D** Table of all detected Crat lysine acetylation sites. Presence/absence in Uniprot or PSP database listed as well as known human acetyl-lysine site or other human PTMs known.

Carnitine O-acetyltransferase (Crat) catalyzes the reversible transfer of an acetyl group between CoA and carnitine. Crat therefore regulates the acetyl-CoA/CoA ratio within the mitochondria with direct implications for mitochondrial carbon flux and metabolic flexibility^66^. Our atlas provides evidence for 18 acetyl-lysine sites on Crat, only three of which appear in either the PSP or UniProt databases (**Figure 5B,D**). Notably none of these sites are directly adjacent to the catalytic histidine, H343; however, one of the novel sites, acK423, resides on a residue that is essential for CoA stabilization^66^. **Figure 5B** presents an illustration of acetyl-CoA binding and a depiction of the electrostatic interaction with the primary amine of K423. Previously, studies have found that despite Crat being expressed across tissues, its activity is primarily elevated muscle, heart, and brown adipose tissue with minimal activity in liver^67^. Interestingly, abundances of acK423 and other acetyl-lysine sites on Crat, mirror these activity profiles, suggesting that lysine acetylation of Crat plays a regulatory role in its activity (**Figure 5C**). This example highlights the hypothesis-generating potential of our lysine acetylation resource for discovery of the metabolic consequences of lysine acetylation.

### Mapping the mouse acetylome to human pathogenic variants highlight clinically relevant sites

To explore potential functional relevance of site-specific lysine acetylation, we leveraged natural amino acid variation, *i.e.*, single nucleotide polymorphisms (SNPs). To accomplish this, we mapped our acetyl-lysine site atlas to the human-to-mouse (H2M)^68^ database to connect mouse sequences with human sequences and their disease relevance (**Supplementary Data 3**). **Figure 6A** summarizes how our acetyl-lysine sites map to amino acid variants, with approximately 1000 non-pathogenic variants and approximately 750 pathogenic variants annotated. Our supposition is that pathogenic variants likely point to functionally relevant lysine residues. Interestingly, we found that highly-abundant acetyl-lysine sites were enriched with more pathogenic-variant residues (**Figure 6B**). Among the over four thousand proteins detected with an acetyl-lysine site, 202 were found to have acetyl-lysines that mapped to pathogenic clinical variants (**Figure 6C**). The number of pathogenic variants on each protein ranged from just one site for 121 proteins, to a maximum of 17 variant sites on the protein isocitrate dehydrogenase 2 (Idh2), followed by 14 sites on fumarate hydratase (Fh) and 13 sites on Idh1.

**Figure 6.**
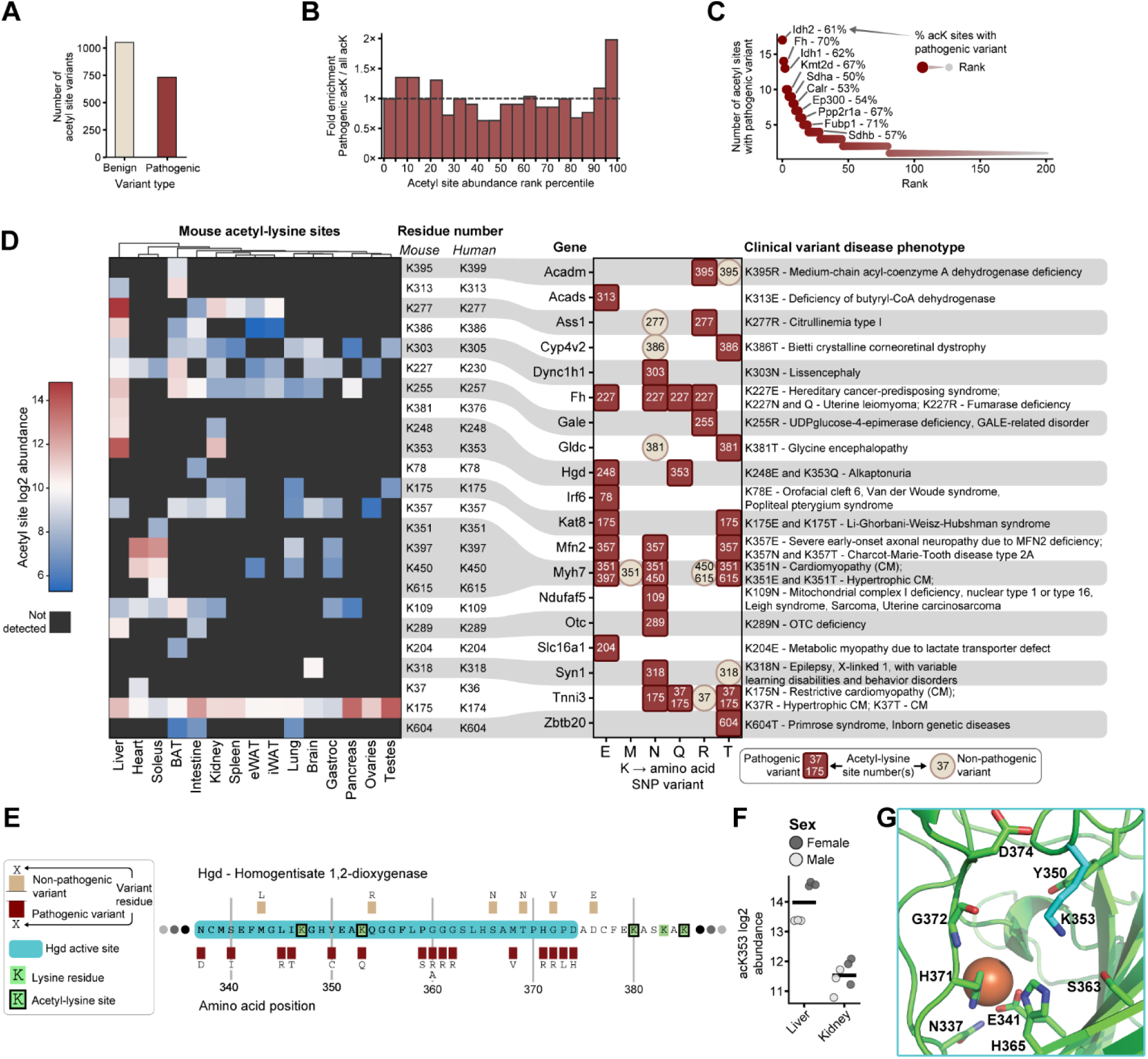
Human clinical variants mapped to mouse acetyl-sites. **A** Number of single nucleotide polymorphism (SNP) clinical variants annotated across all detected acetyl-lysine sites, mapped from the human-to-mouse (H2M) database. **B** Fold enrichment of pathogenic sites compared to background distribution of all acetyl sites. The enrichment is calculated from a background of ranked acetyl site mean abundances. **C** All proteins with detected pathogenic amino acid variants, ranked by the number of acetyl-lysine sites with clinical variants. Percent indicates the proportion of acK sites that were annotated with a pathogenic clinical variant. **D** Summary of clinical variants at acetyl-lysine sites not previously annotated in PSP database. All novel acetyl-lysine sites are given for each protein along with the disease phenotype. The equivalent human protein lysine site number is given, and the acetyl-lysine site abundance from this study is shown. Note that Tnni3 acK175 abundances shown are from a short peptide also mapped to 19 other proteins. **E** Partial sequence of mouse Hgd with active site highlighted (blue) and all reported SNPs with pathogenicity. **F** Abundance of Hgd acK353 in liver and kidney, the only two tissues it was detected in. **G** Structure of Hgd active site from a predicted mouse Hgd structure (EMBL:EBI AF-O09173-F1-v6) overlaid with iron ion position from a human Hgd crystal structure (RSCB PDB ID: 1EY2). Detected lysine acetylation site carbons colored in cyan. Hgd colored in green. Active site substrate atoms colored as follows; carbon: green, nitrogen: blue, hydrogen: red, iron: orange.

We further filtered these results for novel sites not previously annotated in the PSP database and plotted each SNP against the quantitation of its corresponding mouse acetyl-site as well as the amino acid coded for by the mutated variant (**Figure 6D**). We suppose that SNPs that code for a lysine (K) to glutamine (Q) mutation are especially relevant as Q mimics acetyl-lysine residues by removing the positive charge of the lysine at physiological pH while retaining similar chemical properties of the acetyl-lysine side chain^72^.

Focusing on K to Q mutations, we observe three pathogenic variants: fumarate hydratase (Fh) K227Q, homogentisate 1,2-dioxygenase (Hgd) K353Q, and troponin I cardiac (Tnni3) K37Q. We observed four acetyl-lysine sites on Tnni3, one of the three subunits of the troponin complex necessary for heart muscle contraction. Mutations in K37 are associated with hypertrophic and dilated cardiomyopathies (**Supplementary Figure 6A, B**). Notably, K37Q results in dilated cardiomyopathy, while K37R is non-pathogenic^73^. These data suggest that the presence of a positive charge in position 37 of Tnni3 is crucial to its function, and acetylation, similar to a K to Q mutation, would neutralize this charge. In contrast, we observed no pathogenic mutations at the K107 and K118 acetyl-sites, suggesting acetylation is less detrimental at these positions (**Supplementary Figure 6B**). As Tnni3 is cardiac-specific, it was no surprise that we observed these acetyl-lysine sites primarily in heart tissue (**Supplementary Figure 6C**). K37 is found within the tropomyosin complex binding domain of Tnni3^74,75^ (**Supplementary Figure 6D**) and removal of its positive charge via acetylation could negatively impact complex assembly.

Another example of a pathogenic K to Q mutation is found on Hgd, an enzyme that catalyzes the conversion of homogentisate to 4-maleylacetoacetate in the pathway of aromatic amino acid breakdown, whose products feed into ketogenesis. Dysfunction of Hgd leads to alkaptonuria, a condition where the body accumulates homogentisic acid in the blood and tissues. Importantly, Hgd is highly conserved between human and mouse. In our atlas, we detected eleven sites of lysine acetylation on this protein. Two of the lysine residues align with pathogenic SNPs, of which one, K353Q, is located within the active site (**Figure 6E**). Acetylated K353 was only detected in liver and kidney (**Figure 6F**). Crystal structures of human Hgd exist^76^ but do not provide sufficient density in the loop that contains K353; however, a mouse Alphafold predicted structure, which aligns strongly with the human crystal structure, positions K353 at the mouth of the active site (**Figure 6G and Supplemental Figure 6D**).

This region of high flexibility suggests a potential role for K353 in the stabilization of the iron-dependent active site and/or substrate access as previously suggested^73^. Loss of a positive charge due to acetylation, or K to Q mutation, could affect stabilizing interactions with Y347, S363, G372, or D374. We additionally observe disease-associated SNPs for Y347, G372, and D374, suggesting the importance of these interactions for the functional role of Hgd. Therefore, acetylation of K353 in sufficient quantity could either serve a regulatory role or be detrimental towards Hgd function and general organismal health.

## Discussion

In this work we describe the most comprehensive mouse acetyl-lysine atlas. Our efforts leveraged high MS/MS scan rate and sensitivity of the latest generation of Orbitrap Astral hybrid mass spectrometer paired with improved acetyl-lysine enrichment capabilities via more efficient immunoaffinity-based magnetic bead approaches. Together, this combination of technologies afforded identification of ∼18,000 acetyl-lysine sites in under two days of analysis. This resource supports numerous opportunities to explore the role of lysine acetylation across the biological and metabolic space. We utilized these data to investigate lysine acetylation on ribosomes, metabolic enzymes, and human disease-relevant sites.

Globally we observe clustering of acetyl-lysine profiles by tissue primarily driven by shared function. Yet, despite tissue specificity for clustering, we observe nearly 14% shared acetyl-lysine sites across all tissues. This contrasts with prior studies, which observed relative sparsity of acetyl sites across tissues^8^. We believe that this improvement in acetyl site coverage can be attributed to the technical advances leading to higher sensitivity and depth. Our global analysis supports that the lysine acetylome is enriched with mitochondrial proteins. We also corroborated previous reports detailing the affinity for acetylated lysines to be near other lysine residues as well as upstream negatively charged residues. Integrating three-dimensional structural information, we observed the preference of lysine acetylation towards highly structured, less solvent accessible positions within proteins. Whether this result is driven by a true preference for this localization, or through a “survivorship bias” against deacetylation machinery remains to be seen.

Our enrichment analysis highlighted translational proteins as enriched for lysine acetylation. We specifically observed many ribosomal lysine acetylation sites in the liver and pancreas. Notably, these sites are well-conserved between mice and humans and are known ubiquitination sites. Therefore, acetylation of ribosomal proteins could serve to extend ribosomal half-life, whereby acetylation of lysine would prohibit ubiquitin attachment and therefore decrease the probability of proteasomal degradation^57^. This mechanism could provide an avenue for metabolic perturbations, i.e. glucose-load, to sustain ribosomal translation. Our hypothesis is particularly compelling in the context of the pancreas where time-sensitive insulin production necessitates higher translational capacity^77,78^.

We further utilized this resource for analysis of mitochondrial protein acetylation, which was elevated in BAT, liver, and kidney. Protein acetylation in the mitochondria is primarily driven by a non-enzymatic mechanism due to the high localized concentration of acetyl-CoA^48^. One mechanism to maintain acetyl-CoA/CoA homeostasis is the carnitine O-acetyltransferase enzyme, Crat^79,80^. Our atlas identifiesfifteen previously unreported lysine acetylation sites on Crat. We focused on K423, a residue known to stabilize CoA binding within the active site^66^. Crat activity is uniquely upregulated in skeletal and heart muscle^67^; these tissues in turn have the highest abundance of acK423. Together, this suggests a regulatory function of acetylation of Crat K423, highlighting the hypothesis-generating power of our atlas.

To further interrogate functional roles of lysine acetylation, we leveraged the natural variation in human genomes by integrating the H2M database^68^. We hypothesized that disease-associated SNPs could direct focus to lysine residues of functional importance and connect lysine acetylation to human disease. This approach mapped over 700 human pathogenic SNPs to our acetyl-lysine atlas. Among these pathogenic SNPs, we focused on the acetyl-lysine mimicking K to Q mutation. We report three novel acetyl-lysine sites with likely functional and potentially pathogenic consequences: Fh acK227, Hgd acK353, and Tnni3 acK37. This atlas not only describes an inventory of pan-tissue acetyl-lysine profiles but links these sites to clinically relevant variants.

While we have now created the largest mouse lysine acetyl-atlas to date, we expect further improvements as technologies advance. As LC-MS instrumentation continues to improve scan rates, limit of detection, and throughput, this will enable expansion of the known lysine acetylome. Similarly, we anticipate further progress in PTM enrichment techniques. Due to the high likelihood of acetyl-lysine residues to be positioned near other lysine residues, the use of alternative proteases that do not cleave after lysine could expand the number of detectable sites. Lastly, we envision a multi-PTM atlas that could place these acetyl-lysine sites in context with other modifications.

## Methods

### Mouse tissue preparation

All mouse husbandry, tissue harvesting, and related methods were performed in accordance with the National Institute of Health Guide for the Care and Use of Laboratory Animals and were approved by the Animal Care and Use Committee at the University of Wisconsin-Madison. Mice were kept within a humidity range of 30-50% in a 12-hour light/dark cycle at 23°C. Mice were provided with a standard chow diet (Teklad #2018). 47-day old male and female C57BL6/J wildtype mice (n=3 per sex) were euthanized via cervical dislocation and tissues immediately harvested and flash frozen in liquid nitrogen to be stored at −80°C. For each tissue, samples from three mice of the same sex were cryo-pulverized together into powder under liquid-nitrogen and kept on dry ice until aliquoted. For each tissue, ∼30mg frozen powder was resuspended in ∼1.5 mL lysis buffer (6M Urea Sigma-Aldrich AM9856, 100mMTRIS, pH 8 Thermo Scientific 327360010, 10mM TCEP Sigma-Aldrich C4706, 40mM 2-chloroacetamide Sigma-Aldrich C0267), vortexed, and bath sonicated for two sets of 5 minutes at 4°C with 3-minute breaks on ice. Protein concentration was estimated with a protein BCA kit (Pierce, 23235). 4 mg of protein was transferred to a new tube and resuspended in methanol (Optima LC/MS grade, Fisher Scientific) to 90% (v/v) to precipitate protein. Samples were then centrifuged at 9000xg for 5 minutes, then decanted to dry pellet. Samples were resuspended with 0.3 mL lysis buffer (6M Urea Sigma-Aldrich AM9856, 100mMTRIS, pH 8 Thermo Scientific 327360010, 10mM TCEP Sigma-Aldrich C4706, 40mM 2-chloroacetamide Sigma-Aldrich C0267). Protein concentration estimated via Nanodrop (Thermo Scientific). Lysate diluted with 0.9 mL 50 mM TRIS pH 8 (Thermo Scientific 327360010) prior to digestion.

### Protein digestion

To each sample, Sequencing Grade Modified Trypsin (Promega V5113) and Lysyl Endopeptidase (LysC, Wako 129-02541) were added at an enzyme: protein ratio of 1:50 and 1:100 respectively and digested overnight at room temperature (25 °C). Each sample was then acidified with 50 µL of 10% trifluoroacetic acid (Sigma-Aldrich, HPLC grade, >99.9%) to quench the digestion. Samples were then desalted with a Strata-X 33 µm polymeric reversed phase SPE cartridge (Phenomenex, 8B-S100-UBL). Peptides were then dried via a SpeedVac (Thermo Scientific) and stored at −80 °C until following steps.

### Acetylpeptide enrichment

Acetylpeptides were enriched from digested samples using the PTMScan®HS Acetyl-Lysine Motif (Ac-K) Kit(Cell Signaling Technology, 46784) using step III of the CST IAP protocol. Approximately 1 mg of dry peptides were resuspended in 1.5 mL HS IAP Bind Buffer (Cell Signaling Technology, 18494). Solution cleared by centrifugation at 10,000xg for 5 minutes at 4°C. Samples kept on ice until enrichment. Antibody-bead slurry was pulse spun then 20 µL gently aliquoted into a separate tube, one per sample. Magnetic beads were washed on a magnetic separation rack with 1mL ice cold 1x PBS (Cell Signaling Technology, 9808) four times. Peptide solution was transferred to magnetic bead tube and incubated with head-over-head rotation at 4°C for 2 hours. Tube pulse spun and set on magnetic separation rack for 10 seconds. Beads with bound peptide were washed four times with ice cold HS IAP Wash Buffer (Cell Signaling Technology, 18494). Beads then washed twice with ice cold LCMS grade water. Acetylpeptides eluted with two washes/incubation steps of 50 µL IAP Elution Buffer (0.15% TFA, ThermoFisher Scientific, 28904). Buffer incubated with beads for 10 minutes at room temperature and gentle mixing. Enriched acetylpeptides were then desalted with a Sep-Pak Vac 1cc cartridge (50mg tC18, Waters WAT054960) then dried down again via SpeedVac (Thermo Scientific). Each sample was kept at −80 °C until resuspended in 20 µL 0.2% formic acid (Fisher Scientific, LC-MS grade) before peptide concentration estimation via Nanodrop (Thermo Scientific) followed by LC-MS analysis as described below.

### Liquid chromatography-mass spectrometry analysis

Nanoflow capillary columns (75 μm I.D., 360 μm O.D.) with pulled nanoESI emitters were packed to 40 cm using high pressures with C18 particles with a 1.7 μm diameter, 130 Å pore size BEH C18 particles (Waters) as previously described^81^. Samples were separated then analyzed with a Vanquish Neo UHPLC (Thermo Scientific) paired with an Orbitrap Astral Zoom mass spectrometer (Thermo Scientific). Source voltage was set to 2000 V. Mobile phase A and B were 0.2% formic acid in water (Fisher Scientific, Optima LC-MS grade) and 0.2% formic acid/80% acetonitrile (Fisher Scientific, Optima LC-MS grade), respectively. Column temperature was held at 50 °C with a custom-built column heater^82^ with a constant flow rate of 300 nL/min. 250 ng of each sample was loaded for each run with injection triplicates, with some samples being less due to enrichment yield limitations, with a post-processing correction provided by input normalization via the nonenriched runs injected at 250ng/sample as described below. Initial conditions of 0%B were increased to 12% over the course of 0 to 2 minutes. The active gradient spanned 12% to 57%B for 30 minutes with curve type 5 beginning at 2 minutes. Column was washed with 100%B for 5 minutes at the end of each gradient.

MS^1^ spectra were acquired in positive mode with a scan range of *m/z* 380-980 and were collected in the Orbitrap every 0.6 s at a resolving power of 240,000 at m/z 200. MS^1^ normalized AGC target was 300% (3e6 charges) with a maximum injection time of 10 ms and RF Lens was set to 40%. DIA MS^2^ scans were acquired in positive mode in the Astral analyzer over a 140-2000 m/z range with a normalized AGC target of 500% (5e4 charges), a maximum injection time of 3.5 ms, and an RF Lens setting of 50%. An HCD collision energy setting of 27% and a default charge state of +2 were set. Pre-accumulation and window placement optimization was turned on. Isolation width was set at 2 Th with no overlap.

### MS data processing

DIA data was processed in Spectronaut version 20.1.250624 via a library-free search in directDIA mode. The enriched acetylproteomics data was searched against the mouse proteome downloaded from Uniprot from Swiss-Prot on 7.2.2025 and input normalized against nonenriched samples in Spectronaut. Unless specified otherwise, default Spectronaut search parameters were used with variable lysine acetylation modifications included. Identification cutoffs for all precursor and protein Qvalues and PEP values set to 0.01. Spectronaut reports were filtered for 0.01 PG.Qvalue and EG.Qvalue cutoffs. The PTMSiteReport file was used for analysis of lysine acetylation sites, with acetyl sites filtered by rows containing “Acetyl (K)” in the “PTM.ModificationTitle” column and values greater than 0.75 in the “PTM.SiteProbability” column. Unique acetyl sites were identified by filtering for unique values of “PTM.CollapseKey”.

### Processed data filtering and normalization

Outputs from the Spectronaut processing of acetyl peptides and total proteomics were passed through custom Python scripts to filter out low confidence results and normalize across batches. Both datasets underwent a similar set of steps, as follows. First, an acetyl peptide or protein was identified if they were found in at least two replicates in at least one tissue in at least one sex. After log2 transformation, the datasets underwent median normalization at the tissue level. Next, the datasets underwent imputation using the standard Perseus^83^ method, operating feature-wise, with an abundance distribution downshift of 1.8 log2 units and 0.3 distribution scale width multiplier. Finally, gene ontology (GO) term annotations for each protein and acetyl peptide were downloaded from UniProt (accessed 6 Nov. 2025), and the complete GO term hierarchy upwards from the given GO terms was populated for each entry using custom scripting with package goatools (v1.5.1).

### Overview of acetyl sites heatmap

Data for the clustered heatmap was generated by taking the mean of detected values within each tissue, within each sex. Next, the missing values were filled using the imputed dataset, after taking the mean within each tissue, within each sex. To prepare for clustering, the data were standardized within each protein, and clustering was performed using Euclidean distance metric and linkage method average with scipy (v1.11.1). Data were plotted using the package seaborn (v0.13.2) clustermap method, and missing values were masked with gray for the final visual. The acetyl site hierarchical clustering underwent dendrogram cutting using scipy cut_tree method to generate 400 groups. These groups underwent overrepresentation analysis using Fisher’s exact test on GO terms, comparing each group versus all other groups, with p<0.05 significance cutoff.

### PCA and loadings plot

Imputed acetyl site abundance data underwent feature-wise standardization, followed by principal component analysis (PCA) using scikit-learn (v1.7.1). Loadings for all acetyl sites were plotted for PC1 and PC2, and top acetyl sites were pulled out based on highest absolute value of loading in both PC1 and PC2.

### GO term selection for heatmap and unique terms in each tissue

GO term enrichment of acetyl proteins in each tissue was calculated using Fisher’s exact test with significance threshold of p<0.05. When a GO term’s enrichment test could not be calculated in a tissue due to no acetyl sites being found, the p-value was given as 1.0. Next, the p-values were transformed into -log10 p-values, and statistics for the mean and standard deviation of the -log10 p-values was calculated across tissues. GO terms found in the heatmap were selected based on filtering criteria. To be retained, a GO term had to have standard deviation across tissues greater than 0.6, have any tissue with -log10 p-value less than 0.6, and have any tissue with -log10 p-value greater than 3. To remove similar GO terms with many overlapping terms, the filtered list of GO terms was pruned to remove GO terms that had more than 90% overlap with another term. Finally, the filtered list of GO terms was pruned again to enhance specificity of the terms by selecting the most specific descendent GO term if there were any parent-child relationships in the GO terms.

To determine the top 10 most unique GO terms within each tissue, the -log10 p-values within each GO term across tissues were Z-scored, and the GO terms were ranked according to having the greatest difference between that tissue’s Z-score and the mean Z-score across the other 14 tissues.

### Sequence logo plot generation

For sequence logo generation, the flanking regions of acetylation sites were obtained from the ‘PTM.FlankingRegion’ column in the Spectronaut output, yielding residues ±7 residues around the acetylation site. For the sequence logo plot for all lysines in the mouse proteome, the mouse proteome database (downloaded from UniProt^84^ on 7.2.2025) was parsed for all lysine residues. For gene ontology-specific logo plots, the acetylation sites were filtered based on gene ontology IDs obtained from QuickGO^85^ and GO terms assigned to proteins in the UniProt^84^ database. Sequence logo plots were then generated in Python using LogoMaker^86^.

### Structural calculations

The predicted structures for the mouse proteome were downloaded from the AlphaFold Protein Structure Database^87,88^. Each predicted structure .pdb file was parsed with the ‘Bio.PDB’ module within Biopython^50,51^. The predicted local distance different test (pLDDT) values generated by AlphaFold at each residue were obtained from the B-factor fields within the PDB files. The solvent accessible surface area (SASA) values were calculated using the ShrakeRupley algorithm within ‘Bio.PDB.SASA’^49–51^. Gene ontology specific plots were obtained by filtering sites as described above.

For the ribosome-specific calculations, SASA values were calculated based on the experimental structure RCSB PDB: 7CPU.

### Structural imaging

The PyMOL Molecular Graphics System, Version 3.0 Schrödinger, LLC. was utilized for structure imaging of the mouse ribosome (RCSB PDB: 7CPU), mouse Crat in complex with carnitine and acetyl-CoA (RCSB PDB: 2H3P), human Tnni3 (RCSB PDB: 1J1E), human Hgd (RCSB PDB: 1EY2), and mouse Hgd (EMBL:EBI AF-O09173-F1-v6)

## Resources/Data Availability

All mass spectrometry data and search outputs is deposited in the MassIVE database under accession MSV000100290. Code for data analysis and figure generation is located at https://github.com/coongroup/Mouse-acetyl-lysine-atlas.

## Acknowledgements

We are grateful for support from the National Institutes of Health (grants R35GM118110 to J.J.C. and R35GM150899 to A.G.). R.W.S. is supported by the Morgridge Institute for Research. B.J.A. acknowledges support of the Hematology T32 traineeship T32HL007899. We acknowledge assistance from William Beimers for his work on the initial data analysis efforts, Annie Jen for assistance on figure editing, Salma Abou Elhassan for assistance on figure editing, and Matthew Healy for work on figure design. We acknowledge BioRender as the source for mouse and organ icons in Figure 1.

## Author Contributions

J.H. and A.G. collected mouse tissues. R.W.S. performed mass spectrometry sample preparation and data acquisition. R.W.S. and B.J.A. performed mass spectrometry data processing. R.W.S., B.J.A., N.M.L, M.K., and T.G. performed data analysis. R.W.S., B.J.A., N.M.L., K.A.O., and J.J.C. contributed to the figure content and design. R.W.S., B.J.A., K.A.O, N.M.L., and J.J.C. wrote and edited the manuscript. J.J.C. provided funding. J.J.C. and K.A.O. provided project supervision.

## Additional Information

List of supplemental material:

Supplementary figures 1-6

Supplementary data tables 1-3

Correspondence and request for materials addressed to Joshua Coon at.

---

**Supplementary Figure 1.**
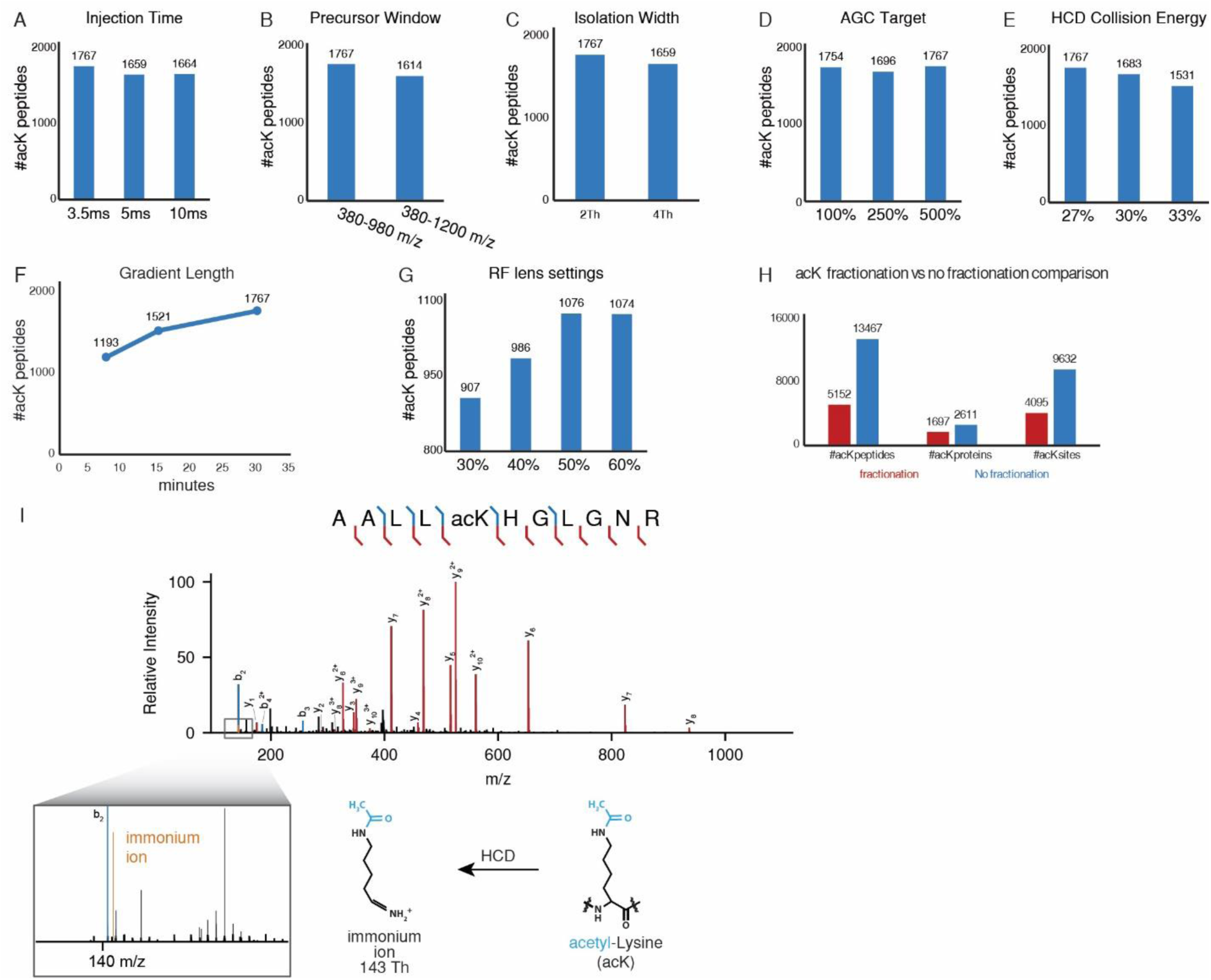
LC-MS parameter sweeping and acetylpeptide protocol optimization. **A-G** LC-MS parameter sweeping. All parameters tested via 250ng male mouse liver acetylpeptide injection with n=1 injection per parameter setting. **H** Comparison between analysis of single batch of enriched acetylpeptides from male mouse liver with our without high-pH fractionation. Fractionation was performed on an Agilent 1260 Infinity BioInert LC with an automated fraction collector over a 20-minute method with a Waters XBridge, Peptide BEH C18, 3.5 µm, 130 Å, 4.6 mm x 150 mm column at 0.8 mL/min as previously described^17^. **I** Tandem mass spectrum of representative acetylpeptide exhibiting inclusion of the lysine acetylation reporter ion, the immonium ion.

**Supplementary Figure 2.**
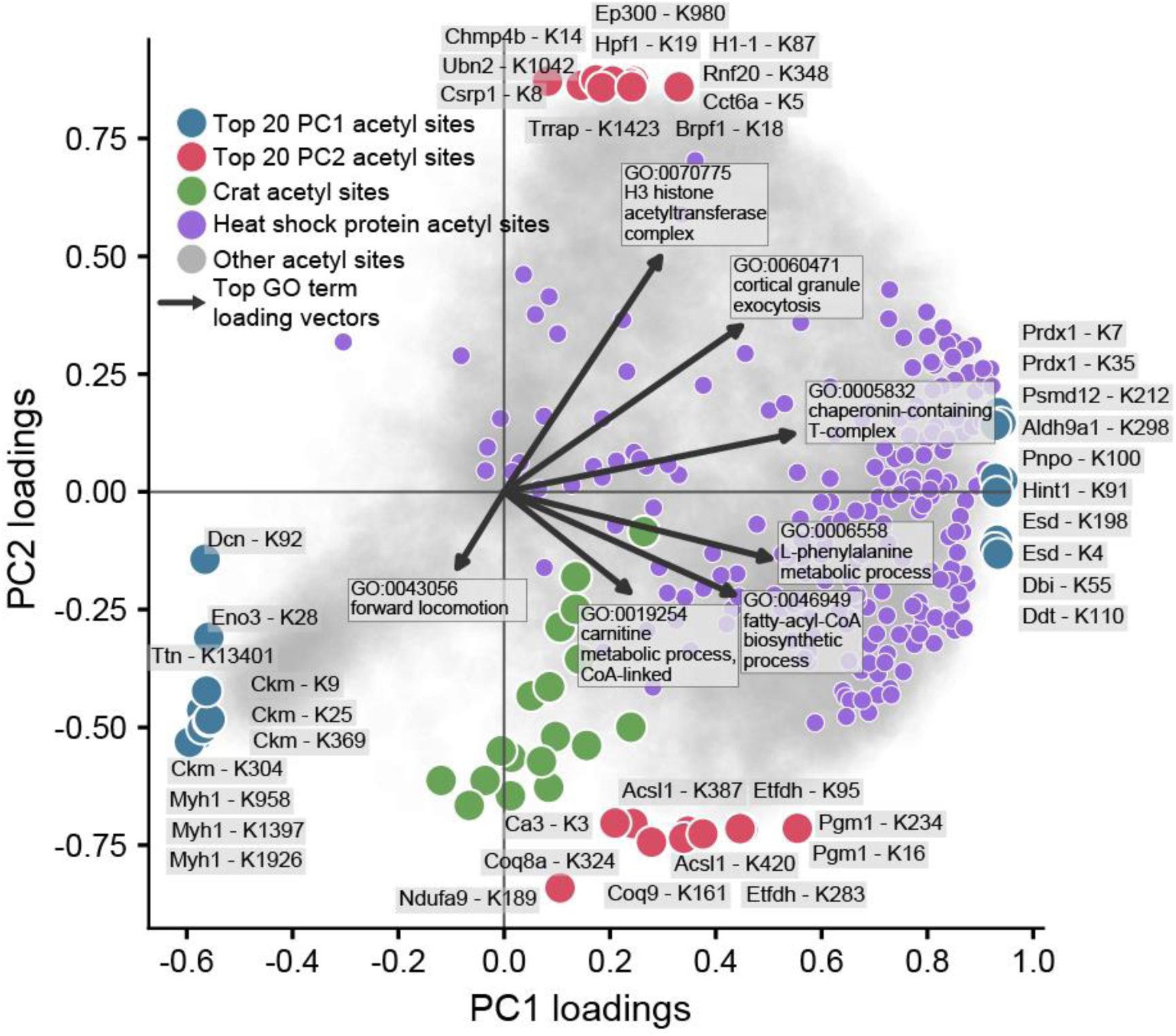
PCA loadings and top loadings drivers. Loadings graph detailing major drivers of Figure 3A PCA.

**Supplementary Figure 3.**
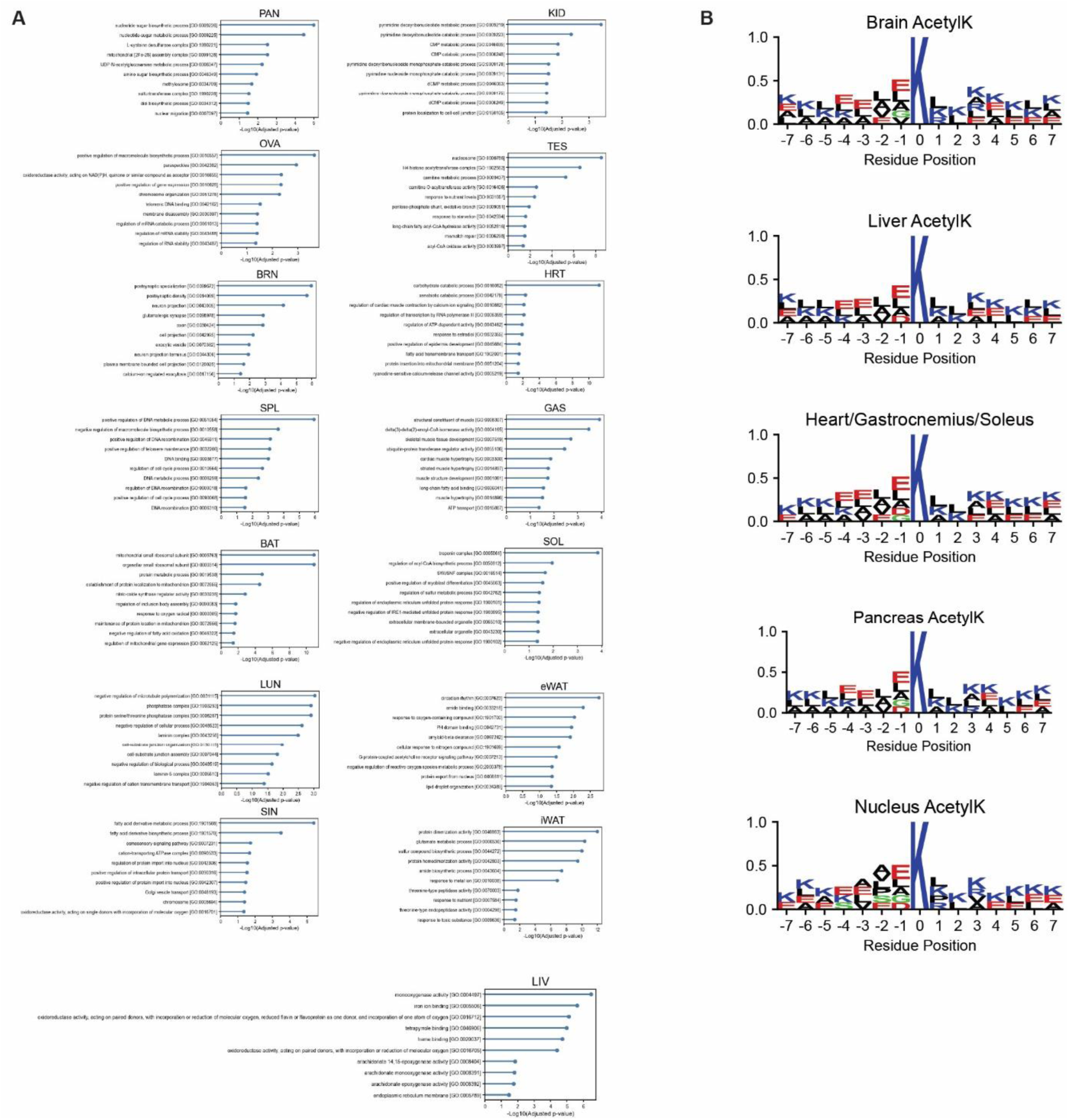
Tissue Specific GO term enrichment and selected sequence motifs. **A** Top 10 GO terms for acetyl-lysine sites found uniquely enriched in each individual tissue. **B** Selected sequence motifs for brain, liver, muscle, and pancreas acetyl sites as well as nuclear-localized acetyl sites.

**Supplementary Figure 4.**
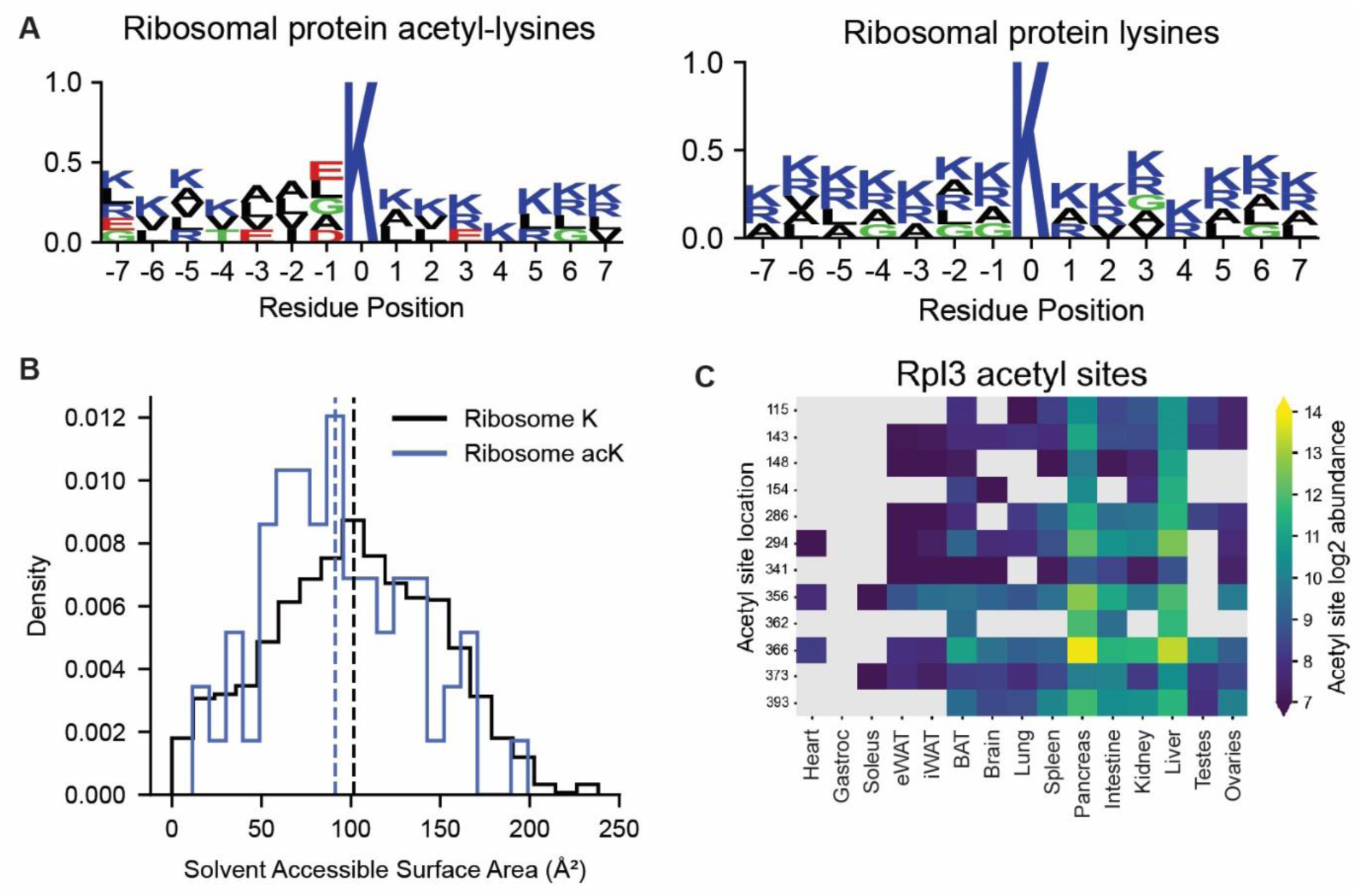
Ribosomal acetylation. Sequence motif of all detected ribosomal protein acetyl-lysine sites. **B** Solvent accessible surface area (SASA) distribution of all ribosomal lysines and all observed ribosomal acetyl-lysines. **C** Clustered heatmap detailing all detected acetyl-lysines found on Rpl3 across all tissues, sex agnostic.

**Supplementary Figure 5.**
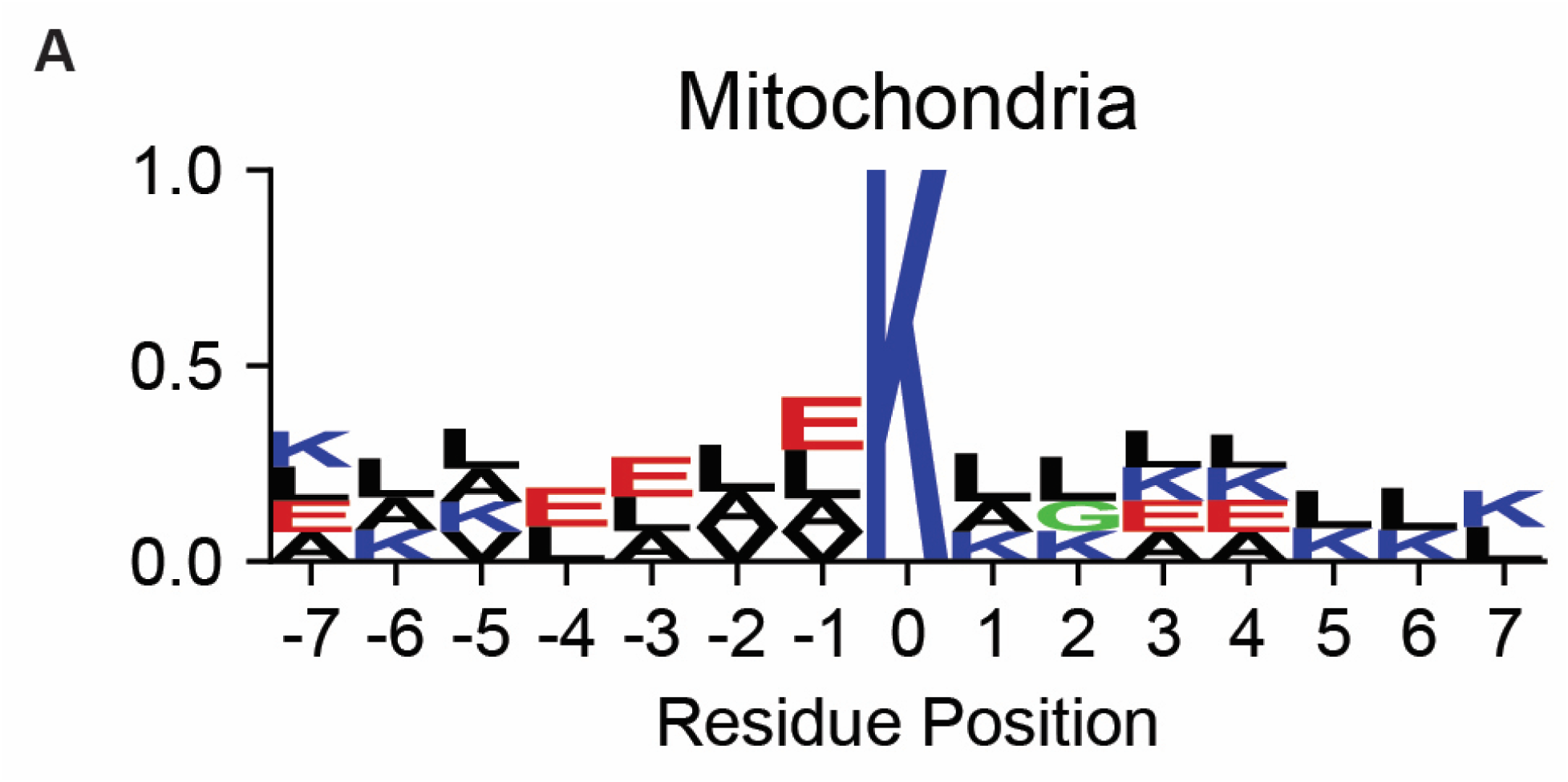
Mitochondrial protein sequence motif.

**Supplementary Figure 6.**
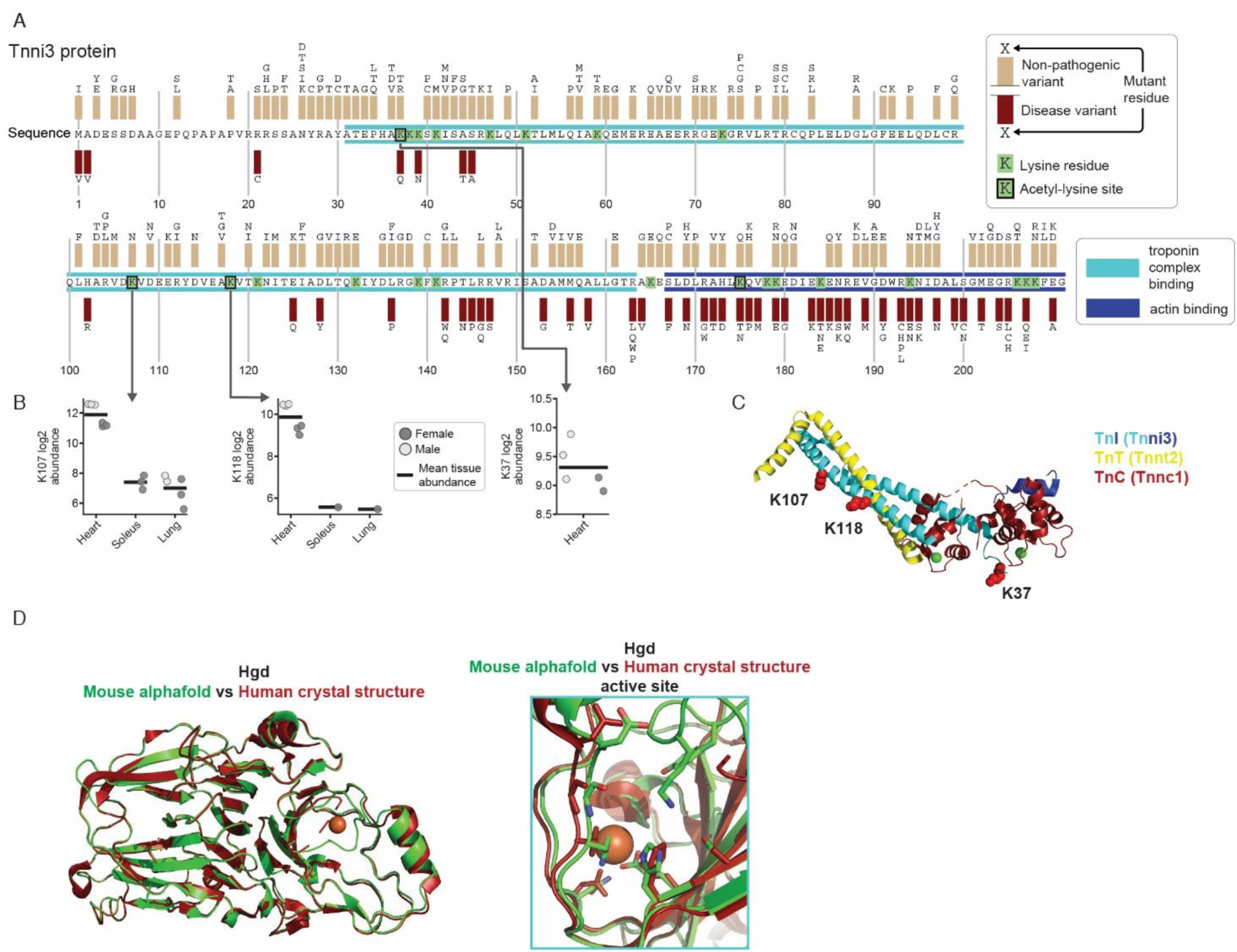
Full H2M acK analysis and Tnni3 acetylation. **A** Sequence of human Tnni3 with troponin complex binding domain highlighted in cyan and the actin binding site highlighted in dark blue. All reported SNPs annotated. **B** Abundance of selected acetyl-sites in mouse tissues. C Structure of human tropomyosin complex crystal structure (RSCB PDB ID: 1J1E). Detected lysine acetylation sites colored in red spheres. Tnni3 colored in cyan and dark blue (same as above). Tnnt2 colored in yellow. Tnnc1 colored in red. D Alignment of red human Hgd crystal structure (RSCB PDB ID: 1EY2) and green mouse Hgd predicted structure (EMBL:EBI AF-O09173-F1-v6) and active site.

**Supplementary Table 1.** Blood glucose levels of mice at time of tissue harvest.

| **Mice** | **blood glucose**<br>**(mg/dL)** |
| --- | --- |
| Male 1 | 174 |
| Male 2 | 216 |
| Male 3 | 183 |
| Female 1 | 144 |
| Female 2 | 145 |
| Female 3 | 169 |

